# Integrated neural dynamics for behavioral decisions and attentional control in the prefrontal cortex

**DOI:** 10.1101/2020.05.06.080325

**Authors:** Yaara Erez, Mikiko Kadohisa, Philippe Petrov, Natasha Sigala, Mark J. Buckley, Makoto Kusunoki, John Duncan

## Abstract

Complex neural dynamics in the prefrontal cortex contribute to context-dependent decisions and attentional competition. To analyze these dynamics, we apply demixed principal component analysis to activity of a primate prefrontal cell sample recorded in a cued target detection task. The results track dynamics of cue and object coding, feeding into movements along a target present-absent decision axis in a low-dimensional subspace of population activity. For a single stimulus, object and cue coding are seen mainly in the contralateral hemisphere. Later, a developing decision code in both hemispheres may reflect interhemispheric communication. With a target in one hemifield and a competing nontarget in the other, each hemisphere initially encodes the contralateral object, but finally, decision coding is dominated by the task-relevant target. These findings further suggest that exchange of information between hemispheres plays a key role when attentional competition resolves. Tracking complex neural events in a low-dimensional activity subspace illuminates integration of neural codes towards task-appropriate behavior, comprising a building block of learned task structure in the prefrontal cortex.

**AUTHOR SUMMARY:** Flexible adaptive processing of information is integral for everyday goal-directed behavior. To unravel dynamic representation of task-relevant information, we analyzed population activity of a primate prefrontal cell sample in a cued target detection task. In a low-dimensional neural subspace, with separate axes for cue, object identity and decision, trajectories showed initial coding of cue and object in the contralateral hemisphere, followed by coding of the behavioral decision across both hemispheres. With target and nontarget stimuli in opposite hemifields, the data suggest initial coding of the contralateral object in each hemisphere. Object coding is then rapidly shut off for the nontarget, and followed by bilateral coding of the target decision. The results bring detailed insight into task structure and information flow within and between the two frontal lobes as a decision is made and attentional competition is resolved.

## INTRODUCTION

Flexible and adaptive processing of information is a hallmark of goal-directed behavior. With a constantly changing environment, varying task demands determine how input information will be processed and selected, with a key role played by the prefrontal cortex (1–5). In non-human primates, responses of individual neurons are tuned to multiple task variables including cues, stimuli, categorical information, and decisions (6–9). Tuning is adjusted by behavioral relevance, with information more strongly encoded when it is relevant to a current decision (6,10–13). Similar flexible and adaptive coding has been observed in the human frontal cortex using neuroimaging, with representation of behaviorally-relevant task events such as cues, stimulus information, categorical distinctions, and responses (14–22).

To organize task performance, multiple variables must be combined into the correct computational structure or control program (23–25). Correct computational structures must be based on the animal’s long-term learning, indicating how environmental and other variables can be combined for goals to be achieved. To analyze such structures, methods are needed to extract multiple variables from prefrontal population activity, and to study dynamics as these variables are appropriately combined (23,26–29).

In a recent study from our group, Kadohisa et al. (26) asked how prefrontal population activity in nonhuman primates develops when items compete for attention. In a cued target detection task, a cue at the start of each trial indicated the target object to be detected in a subsequent choice display. If a target was present in the choice display, the animal awaited a subsequent go signal, then made a saccade to the target location. Importantly, some displays contained two items presented in opposite hemifields, the target on one side and a nontarget on the other. For these displays, measures of population activity showed intriguing dynamics. Initially, responses were dominated by the item presented contralaterally to the recorded hemisphere, with activity resembling the response to the contralateral item presented alone. Other tasks have revealed a similar contralateral dominance in both prefrontal (13) and inferotemporal (30) neurons. Later, however – in the period leading up to the go signal - activity approached response to the most relevant item (target) presented alone. Critically, dominance by the target was achieved more slowly when the nontarget was itself a target on other trials, consistent with long-established behavioral findings showing that such nontargets are hard to ignore (31). In line with other studies of prefrontal cortex (12,32), target-selective responses appeared to reflect an abstract behavioral categorization rather than a prepared motor command, with no evidence of activity time-locked to the saccade finally made to the target location.

While these results provide initial insight into prefrontal dynamics during attentional selection, the neural process underlying competition and decision is potentially much more detailed and rich, reflecting the learned computational task structure. In such a task, cue and object information are combined to form a final decision for each stimulus in the display (27,33). Here, we disentangle coding of cue, object and decision and track their temporal development in single-stimulus trials, unraveling the codes that construct the control program. We use a recently developed dimensionality reduction technique, demixed principal component analysis (dPCA) (28), to construct a task-related lowdimensional subspace using the same cued target detection task data set used by Kadohisa et al. (26). We then use the extracted neuronal weights of this program to reveal how they implement decisionmaking under attentional competition. Our results show how coding of cue, object and decision evolve and interact in the prefrontal cell population. Importantly, the subspace allows us to not only track dynamic shifts in representation across the state space (e.g.., *when* object and decision information emerge), but also to identify *what* was represented for each stimulus. We provide a detailed characterization of computation within the task-related state space of the control program and its extension to competing stimuli. We show how codes of task variables dynamically evolve across the two hemispheres while competition is resolved, and propose a prefrontal mechanism for resolving attentional competition in this task.

## RESULTS

In this task (Figure 1) two objects were associated with one cue each, and therefore could serve as either targets (T) or distractors (D) in the choice display. A third object was not associated with any cue, and was therefore a neutral (N) stimulus that could never be a target. In single-stimulus displays, choice objects appeared either to the right or left of a central fixation dot, i.e., contralateral or ipsilateral to the recorded hemisphere. To test for attentional competition, two-stimulus displays contained one stimulus in each hemifield, with combinations of target and neutral objects (T + N, low competition), target and distractor objects (T + D, high competition), and distractor and neutral objects (D + N, target absent trials). When the target was present, the animal was rewarded for a saccade to its location, made on receipt of a go signal presented shortly after display offset. When the target was absent, reward was given for maintained fixation. We tracked the dynamics of cue, object and decision information within the task-related subspace and how it evolves when two items compete for attention.

**Figure 1:**
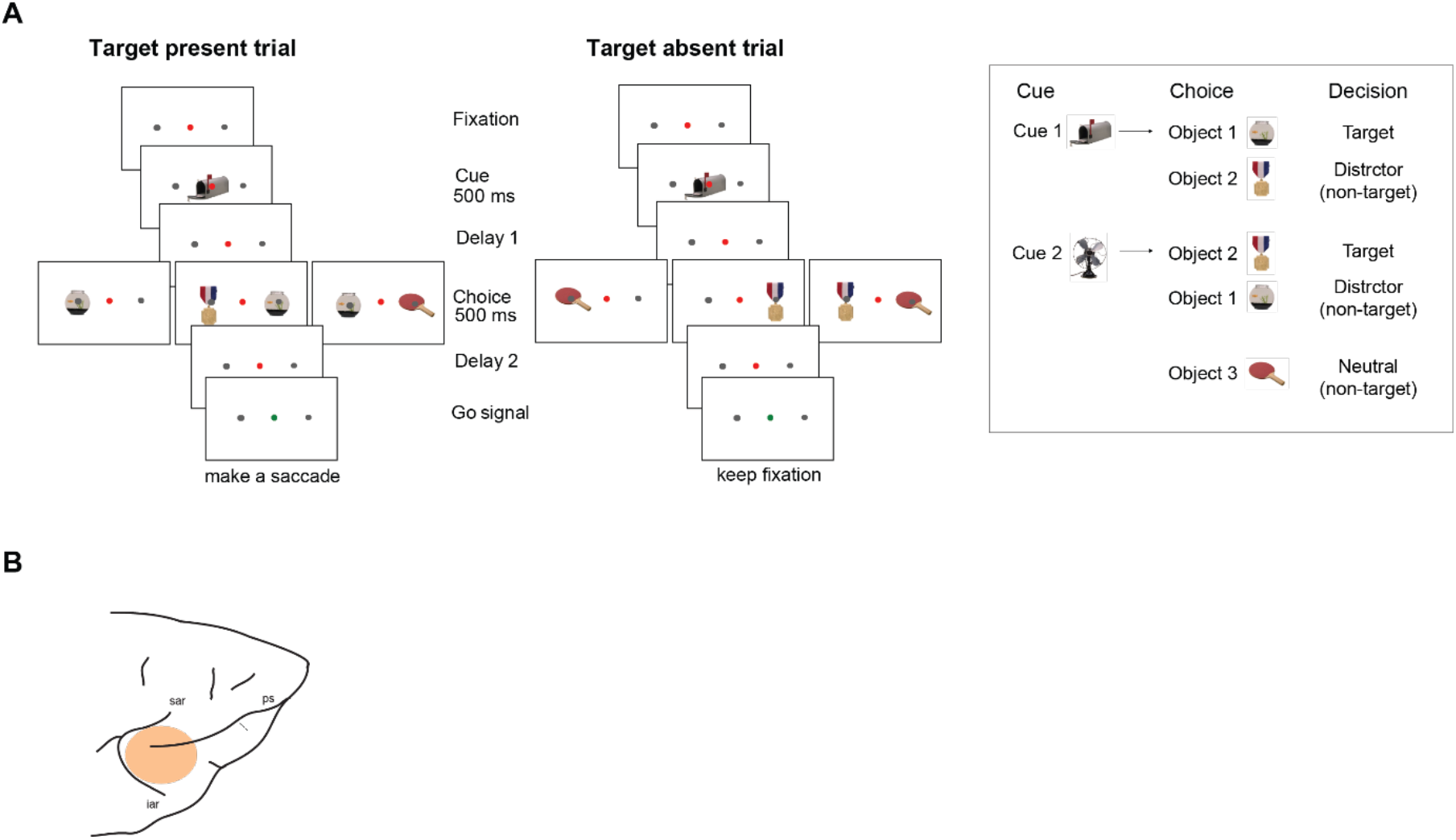
Task and recording locations. **A: Task.** Each trial began with fixation on a central dot, followed by a cue (500 ms). After a delay (variable delay 1, see Materials and Methods for details), a choice display appeared (500 ms) with either one or two stimuli. For each animal, two cues were associated with one target object each based on previous training. Cues and choice objects for one animal are illustrated in the inset on the right. Depending on the preceding cue, the animals had to make a decision whether a target (the associated object) appeared in the choice display (target present trial) or not (target absent trial). Following another delay (variable delay 2, see Materials and Methods for details), the color of the fixation dot was changed to green, indicating the go signal. On target present trials, animals made a saccade to the target location and were then rewarded with a drop of liquid for a successful trial. On target absent trials, animals had to hold fixation during the response interval and were rewarded at the end of a wait period. Choice display objects could be the object associated with the preceding cue (target), the object that was associated with the other cue (distractor), or a third object that was not associated with any cue (neutral). Stimuli could appear either to the right or left of the fixation dot (i.e., contralateral or ipsilateral to the recorded hemisphere). Two-stimulus displays were target + distractor (T + D), target + neutral (T + N) and distractor + neutral (D + N). **B: Approximate recording locations in the right hemisphere.** For one animal, additional recordings were made in a similar region of the left hemisphere. ps: principal sulcus; sar: superior arcuate sulcus; iar: inferior arcuate sulcus.

Performance in the task was overall high, as previously described for this data set (26). For single-stimulus displays, accuracy levels were high for target (average 84%; 86% monkey A, 83% monkey B) and neutral trials (average 85%; 83% monkey A, 87% monkey B). The relatively high difficulty level for distractor trials (which may be targets on other trials) was evident in a much lower accuracy (average 60%; 56% monkey A, 63% monkey B). Most (72%) of the errors in distractor trials were a saccade to the stimulus location following the go signal, further confirming the difficulty of ignoring distractors. Accuracy levels were similar for T + D (average 76%; 70% monkey A, 82% monkey B) and T + N (average 77%; 77% monkey A, 76% monkey B) displays, but while many errors in the T + D displays were saccades to the distractor location (59%), the most common error for T + N displays was to keep fixation (79%). Accuracy levels for the D + N displays were slightly lower (average 67%; 60% monkey A, 73% monkey B), as expected when targets are absent in visual search.

Neurons were recorded during performance of the task on the lateral frontal surface across three hemispheres of the two monkeys. Recording locations (Figure 1B) were in dorsal and ventral regions of the posterior lateral prefrontal cortex, around the posterior third of the principal sulcus. Activity from 461 neurons was recorded, and the analysis included data from 337 neurons that had a minimum of four trials per condition for all conditions (Monkey A: N = 140 right, N = 71 left; monkey B: N = 126 right). The average number of trials per condition was 14.5.

### Heterogeneous responses and mixed selectivity of prefrontal units

Activity of single units was heterogeneous and mixed with diverse selectivity for cue, object and decision. In Figure 2 this is illustrated with responses to different categories of single-stimulus trials for five single units, each tuned to one or more task variables with diverse dynamics. These profiles of activity at the single unit level are in line with previous reports of mixed selectivity of individual prefrontal neurons (6–9). Mixed selectivity and distributed coding of information have been shown to be critical for the wide variety of task-related computations performed in the prefrontal cortex (8).

**Figure 2:**
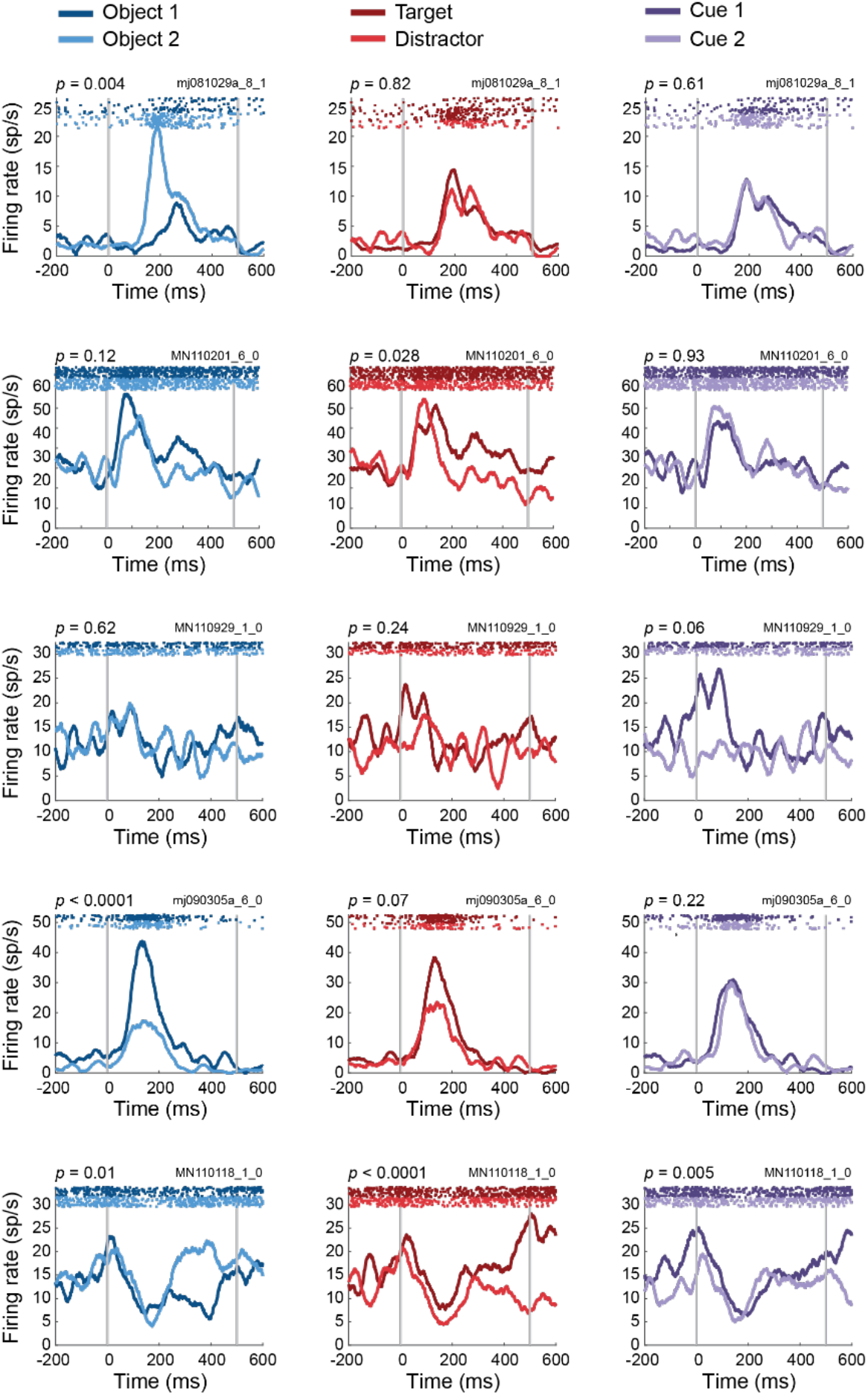
Mixed selectivity and heterogeneous response profiles of single prefrontal cells. Examples of five units (one in each row) and their selectively to object, decision and cue (left, middle and right column, respectively). Data come from single-stimulus trials. Peri-stimulus time histograms show average responses across trials for each of the objects, decisions and cues based on the four single-stimulus contralateral target and distractor conditions. Single trial raster plots are at the top of each panel. Single cells showed a variety of responses to task-relevant variables, some with selectivity to just one variable (object, decision and cue for units shown in 1^st^, 2^nd^ and 3^rd^ rows, respectively), and some with mixed selectivity to more than one variable (object and decision in 4^th^ row, and all three variables in 5^th^ row). Gray vertical lines indicate stimulus onset (0 ms) and offset (500 ms). *p* values above each panel indicate selectivity to each task variable as computed by a two-way ANOVA (main effects for object and decision, interaction for cue, based on trial data averaged across all time points from 200 ms before to 600 ms after display onset). Unit ID is shown above each panel on the right.

### Low-dimensional state space captures coding of task-relevant variables across the neural population

To reveal the underlying task-related representational space, distributed population activity across neurons can be used, unravelling the complex nature of prefrontal cortex activity that implements the learned task structure. Of particular interest is a low-dimensional subspace that can identify population-level representations and their coding of task-relevant variables. To reveal such a subspace, we used demixed PCA (dPCA) (28), a recently developed technique that both decomposes the population activity into components that capture the majority of the variance in the data as well as separates them in respect to task variables such as object and decision. We applied dPCA to construct the low-dimensionality subspace comprised of the three task variables (cue, object identity, and decision), using four single-stimulus conditions that cross these variables: cue1-object1-T, cue2-object2-T, cue2-object1-D, cue1-object2-D. For the dPCA, trial data for these four conditions could be grouped according to object identity (object 1 or 2), decision (T or D) and cue (cue 1 or 2, the interaction of object and decision) to allow the extraction of the relevant components. The analysis focused on the choice display phase of the trial; subspaces were constructed based on trial data from 200 ms before choice display onset until 600 ms after that (100 ms post choice display offset), during which period the animals fixated throughout. Because strong hemifield effects have been previously demonstrated in the prefrontal cortex (26,30,34–37), we constructed separate subspaces for contralateral and ipsilateral stimulus displays. Each hemifield subspace was constructed using a splithalf approach (separation of odd and even trials) to allow for cross-validated data projection (see below). The variance explained by each dPCA component was related primarily to one task variable only, confirming that the contributions of the cue, object and decision were well separated in the subspaces. Each of the first 20 dPCA components was associated with the task variable that explained most of its variance, with some components capturing variable-independent variance driven mostly by the time course of activity during the choice display, not specifically associated with any of the task variables. The first variable-associated component of each of the three task-relevant variables (cue, object and decision) captured a large proportion of the variance related to this variable: it explained on average 2.14 times more variance than the second component associated with this variable and accounted on average for 39% of the variance explained by this variable across the first 20 dPCA components. We therefore constructed, for each hemifield and data-half, a three-dimensional compressed subspace with axes comprised of the first object, decision and cue components. To investigate the temporal trajectory of information coding for each of the task variables, we projected population responses onto each axis of the subspace. To avoid over-fitting, data from each half of the trials were projected onto the subspace of the other half, and projections were averaged across halves.

For the object axis, cross-validated projections for the four critical single-stimulus conditions (cue1-object1-T, cue2-object2-T, cue2-object1-D, cue1-object2-D) are shown in Figure 3A. Top left and bottom right panels show projections of contralateral and ipsilateral stimuli onto the component for their own hemispace, while bottom left and top right panels show cross-projections of each stimulus onto the component extracted for the opposite hemispace. For contralateral stimuli (top left), there was clear separation of object 1 and object 2 for both target and distractor conditions, as expected for a component designed to maximize this separation. Beyond this expected separation, the object component reveals the temporal progression of object information coding, with significant coding starting from around 50 ms post stimulus onset and a peak around 140 ms which then drops and remains sustained throughout the trial. Object coding was much weaker for ipsilateral stimuli (Figure 3A, bottom right), with significant coding from around 100 ms until just after 340 ms, but without the clear peak shown by contralateral stimuli. As expected given relatively little object information for ipsilateral stimuli, both cross-projections (bottom left, top right) also showed only weak separation of the two object identities.

**Figure 3:**
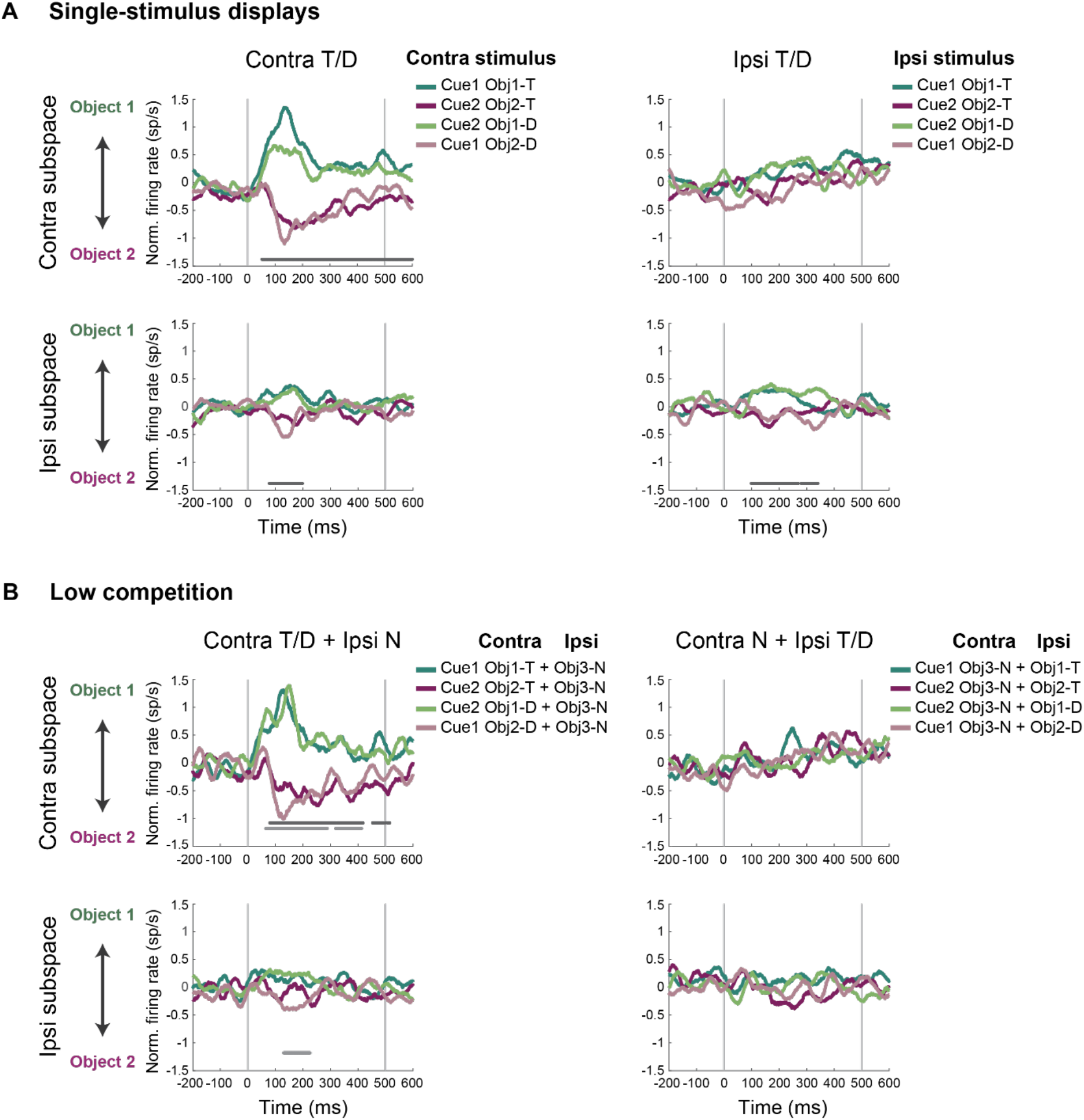
Object information for single-stimulus and low competition conditions. **A**: Projections on the dPCA object axis for single target (T) and distractor (D) stimuli. Population activity relative to choice display onset is projected onto the first object component of each hemifield subspace, with positive firing rates indicating a representation of object 1, and negative firing rates indicating object 2. Projections are plotted for stimuli presented in either contralateral or ipsilateral hemifield (left and right column, respectively), with projections in both the contralateral and ipsilateral subspaces (top and bottom row, respectively). Cross-validated responses are shown as averages of the two halves of the data (odd and even trials), each projected onto the subspace of the other half of the data. Gray vertical lines indicate stimulus onset (0 ms) and offset (500 ms). Horizontal line at the bottom indicates significant difference between object 1 and object 2, averaged across target and distractor. Significance is determined using a permutation approach with cluster-based correction across time points. **B**: Object information for low competition conditions with an added neutral (N) stimulus (T + N and D + N). Horizontal lines at the bottom indicate significant difference between object 1 and object 2 for T + N (dark gray) and D + N (light gray).

For these same single-stimulus displays, a strikingly different temporal trajectory was observed for decision coding (Figure 4A). For both contralateral (top left) and ipsilateral (bottom right) stimuli, projections onto their own subspace showed gradual development of the target-distractor distinction, beginning around 100 ms and 250 ms, respectively, and increasing over time. Cross-projections of each stimulus onto the subspace for the opposite hemifield (bottom left, top right) also showed significant decision information. Responses were weaker than the decision responses in the same-hemifield subspaces, suggesting partial but not complete overlap of decision coding on the two sides.

**Figure 4:**
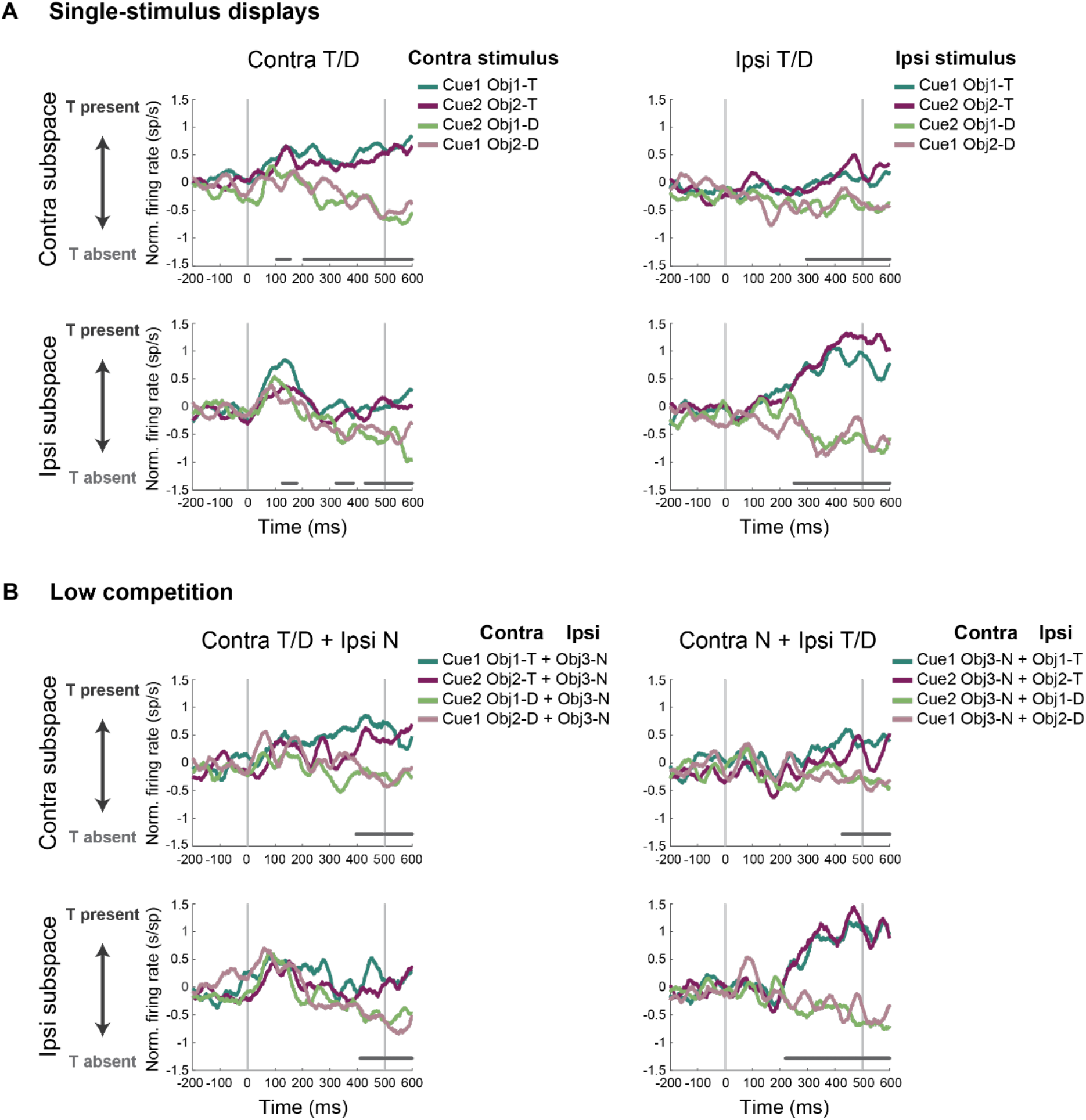
Decision information for single-stimulus and low competition conditions. **A**: Projections on the dPCA decision axis for single target (T) and distractor (D) stimuli. Population activity relative to choice display onset is projected onto the first decision component of each hemifield subspace, with positive firing rates indicating ‘target present’, and negative firing rates indicating ‘target absent’. Projections are plotted for stimuli presented in either contralateral or ipsilateral hemifield (left and right column, respectively), with projections in both the contralateral and ipsilateral subspaces (top and bottom row, respectively). Cross-validated responses are shown as averages of the two halves of the data (odd and even trials), each projected onto the subspace of the other half of the data. Gray vertical lines indicate stimulus onset (0 ms) and offset (500 ms). Horizontal line at the bottom indicates significant difference between ‘target present’ and ‘target absent’, averaged across objects 1 and 2. Significance is determined using a permutation approach with cluster-based correction across time. **B**: Object information for low competition conditions with an added neutral (N) stimulus (T + N and D + N). Gray horizontal line at the bottom indicates significant difference between T + N vs. D + N conditions.

Lastly, projections onto the cue axis revealed some weak cue coding (Figure 5A). Cue information was strongest for a contralateral stimulus projected onto its own subspace, beginning around stimulus onset. As the cue was the same whether the subsequent stimulus was contralateral or ipsilateral, and was known before stimulus onset, a tentative interpretation is reinstatement of a cue signal as a contralateral stimulus is processed. For ipsilateral stimuli, there were weak hints of cue coding in either subspace, though this was only significant for a brief period around stimulus onset, and only for the ipsilateral subspace.

**Figure 5:**
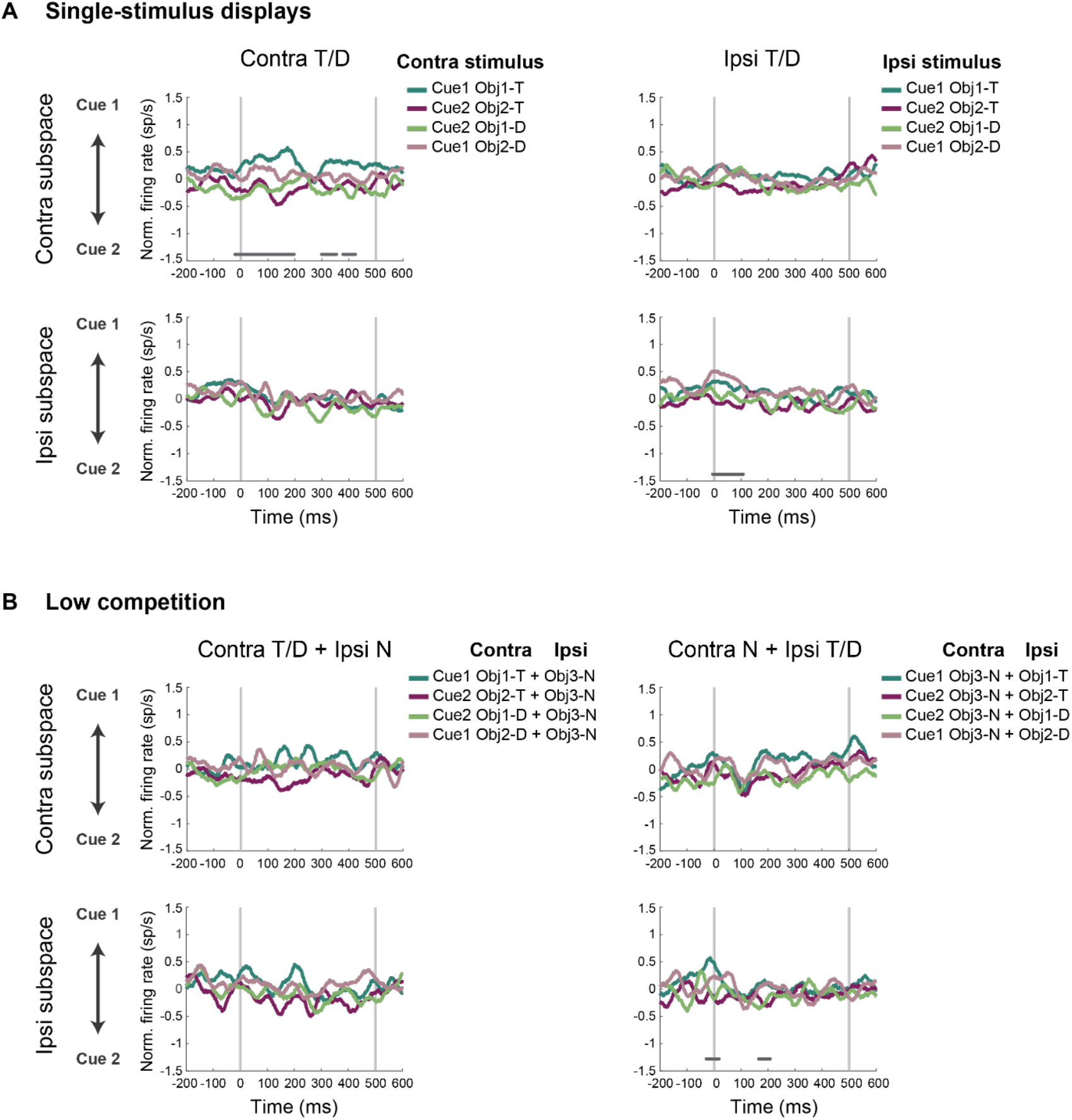
Cue information for single-stimulus and low competition conditions. **A:** Projections on the dPCA cue axis for single target (T) and distractor (D) stimuli. Population activity relative to choice display onset is projected onto the first cue component of each hemifield subspace, with positive firing rates indicating a representation of cue 1, and negative firing rates indicating cue 2. Projections are plotted for stimuli presented in either contralateral or ipsilateral hemifield (left and right column, respectively), with projections in both the contralateral and ipsilateral subspaces (top and bottom row, respectively). Cross-validated responses are shown as averages of the two halves of the data (odd and even trials), each projected onto the subspace of the other half of the data. Gray vertical lines indicate stimulus onset (0 ms) and offset (500 ms). Horizontal line at the bottom indicates significant difference between cue 1 and cue 2, averaged across objects 1 and 2. Significance is determined using a permutation approach with cluster-based correction across time points. **B:** Cue information for low competition conditions with an added neutral (N) stimulus.

Altogether, the single-stimulus data show that task-relevant information is captured by lowdimensional subspaces, with distinct temporal profiles for the representation of the elements that comprise the control program of the task – object, decision and cue. Most prominently, an early phase of strong object coding along with weak cue coding is dominated by the contralateral stimulus. In contrast, a gradual build-up of the decision state is seen for both contralateral and ipsilateral stimuli, with partial overlap of decision coding for stimuli on the two sides.

### Stimulus and decision information in attentional competition

We next used the same subspaces (derived from single-stimulus trials) to investigate contextdependent coding across the neural population when items compete for attention. For comparability with the analyses of single-stimulus data, we again divided trials into two halves, projected data for each half onto the single-stimulus subspace derived from the other half, and averaged the results. Data were analyzed separately for low competition (target/distractor + neutral, T/D + N) and high competition (target + distractor, T + D) displays.

Results for low-competition displays are shown in Figures 3B, 4B and 5B. Comparison with results for a single T/D stimulus (Figures 3A, 4A, 5A) suggests only modest effects of the added N. For object coding (Figure 3B), there was again strong discrimination only for a contralateral stimulus (target or distractor) projected onto the contralateral hemispace, with separate statistical tests for T + N and D + N displays showing similar coding for targets and distractors. In this case, indeed, ipsilateral coding was not significant at any time point (Figure 3B, bottom right). Decision coding was again seen for both contralateral and ipsilateral stimuli (Figure 4B), with a suggestion of delayed onset compared to single-stimulus displays, in particular for a contralateral T/D stimulus on the contralateral subspace. Cue coding again was weak, with only scattered points of significance for an ipsilateral T/D stimulus in the ipsilateral subspace (Figure 5B). Statistical comparisons between single-stimulus and low-competition displays (see Materials and Methods) showed no significant differences in either object, decision, or cue coding, for either contralateral or ipsilateral T/D stimuli, projected onto contralateral or ipsilateral subspaces.

In the high-competition conditions, a target and distractor appeared in the same display (T + D). For these displays, responses on object, decision and cue axes of contralateral and ipsilateral subspaces are shown in Figure 6.

**Figure 6:**
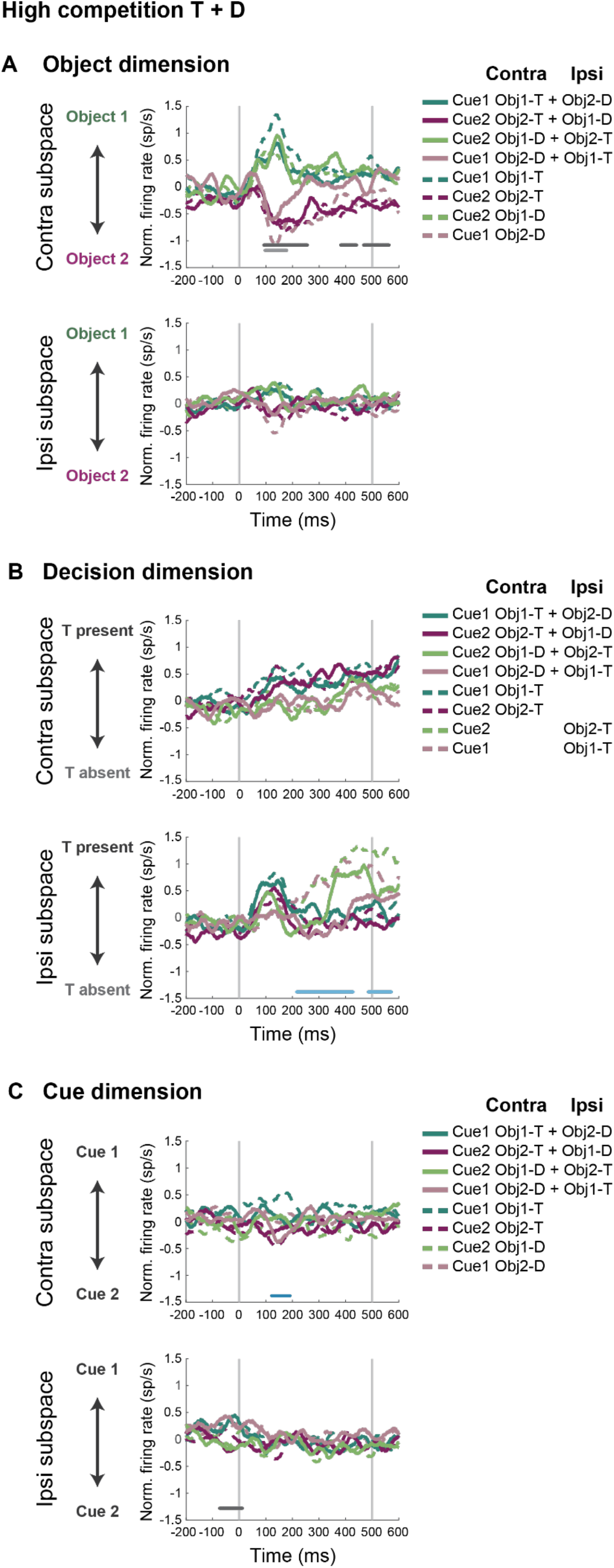
Stimulus, decision and cue coding for high competition conditions (target + distractor, T + D). **A**: Object coding. For comparison, projections of single T/D presented contralaterally (copied from Figure 3A) are plotted as dotted lines. Dark and light gray significance lines at the bottom indicate significant object coding, based on the identity of the contralateral object, for the averaged contra T + ipsi D, and contra D + ipsi T displays, respectively. Statistical comparison between single-stimulus and high-competition displays showed no significant difference in coding the identity of the contralateral object, whether this was T (cyan, purple) or D (green, pink). **B**: Decision coding. For comparison, projections of a single T presented contralaterally or ipsilaterally (copied from Figure 4A) are plotted as dotted lines. There was no difference in decision coding between single-stimulus and high-competition displays when targets were contralateral. Light blue horizontal line at the bottom indicates significant difference in decision coding between single-stimulus and high-competition displays when targets were ipsilateral. **C:** Cue coding. Gray horizontal line at the bottom indicates significant cue coding. There was no difference in cue coding between single- and two-stimulus displays. All other details as in Figures 3–5.

For object coding (Figure 6A), responses to the four possible T + D displays (solid lines) are shown along with responses to a single contralateral stimulus (dotted lines, copied from Figure 3A) for comparison. In the contralateral subspace, responses to T + D resembled responses just to the object in the contralateral field, when this was presented alone. The most striking exception was that, when the contralateral object was a distractor (Figure 6A, solid green and pink), its object-selective response was short-lived, significant only around 100-180 ms from display onset (Figure 6A, pale grey line), suggesting a rapid shut-off of object coding for a contralateral D. Statistical comparison of object discrimination in single-stimulus and T + D displays suggested a brief difference at around 200 ms, significant (*p* < 0.001) for individual time points but not surviving cluster correction. In the ipsilateral subspace there was only a hint of object coding, in line with the weak ipsilateral object coding when these objects were presented alone (Figure 3A).

For decision coding (Figure 6B), responses to T + D displays are shown along with responses to a single contralateral or ipsilateral target (dotted lines, copied from Figure 4A) for comparison. In the contralateral subspace, responses to the T + D display closely followed those for its single target, contralateral or ipsilateral, when this was presented alone. Thus, unlike responses on the object dimension, activity on the decision dimension was driven by the target stimulus, whose presence determined the required response. In the ipsilateral subspace, results were very different. When the T + D display contained an ipsilateral T (Figure 6B, solid green and pink), the activity indicating a ‘target present’ decision was delayed and weakened compared with activity for that same ipsilateral T presented alone (Figure 6B; dotted green and pink; significant difference between single-stimulus and T + D displays shown by pale blue line). Thus population activity indicating an ipsilateral target was significantly suppressed by the presence of a competing, contralateral distractor.

For the cue dimension (Figure 6C), finally, responses on T + D trials again suggested only weak and occasional cue discrimination, resembling that seen in single-stimulus displays (Figure 5A), with a brief difference between single-stimulus and T + D displays on the contralateral subspace around 120-190 ms (Figure 6C, blue line).

Coding along the stimulus and decision dimensions, in which strong coding was observed, is shown as trajectories in the state space in Figure 7. The same projections from Figures 3, 4 and 6 are shown, from choice display onset (black circles) to 600 ms post display onset (red diamonds), for the contralateral (top) and ipsilateral (bottom) subspaces separately. For single-stimulus displays (Figure 7A, same data as in Figures 3A and 4A, top left and bottom right panels), object coding is strong for a contralateral stimulus projected onto the contralateral subspace (top), with an early peak that later decreases, while coding along the decision axis gradually develops. Object coding is weak for an ipsilateral stimulus projected onto the ipsilateral subspace (bottom), with movement along the decision axis similar to the contralateral stimulus/subspace. For low competition (Figure 7B, same data as in Figures 3B and 4B, top left and bottom right panels) trajectories are highly similar to those of the single-stimulus conditions, showing little effect of an added neutral stimulus on coding. Coding when competition is high (Figure 7C, same data as in Figure 6A, 6B) shows large movement along the object dimension in the contralateral subspace (top), dominated by the contralateral stimulus. In this subspace, the decision trajectories all move towards the ‘target present’ state, with coding similar to when the targets are presented alone (contralateral target similar to a single contralateral target as in Figure 7A, top; ipsilateral target similar to a single ipsilateral target as in Figure 4A, not shown in Figure 7A). The interrupted coding in the ipsilateral subspace (bottom) when competition is high is seen in the limited movement in the state space for all displays, in particular along the decision axis for an ipsilateral target (green and pink) compared to a single ipsilateral target (cyan and purple in Figure 7A, bottom).

**Figure 7:**
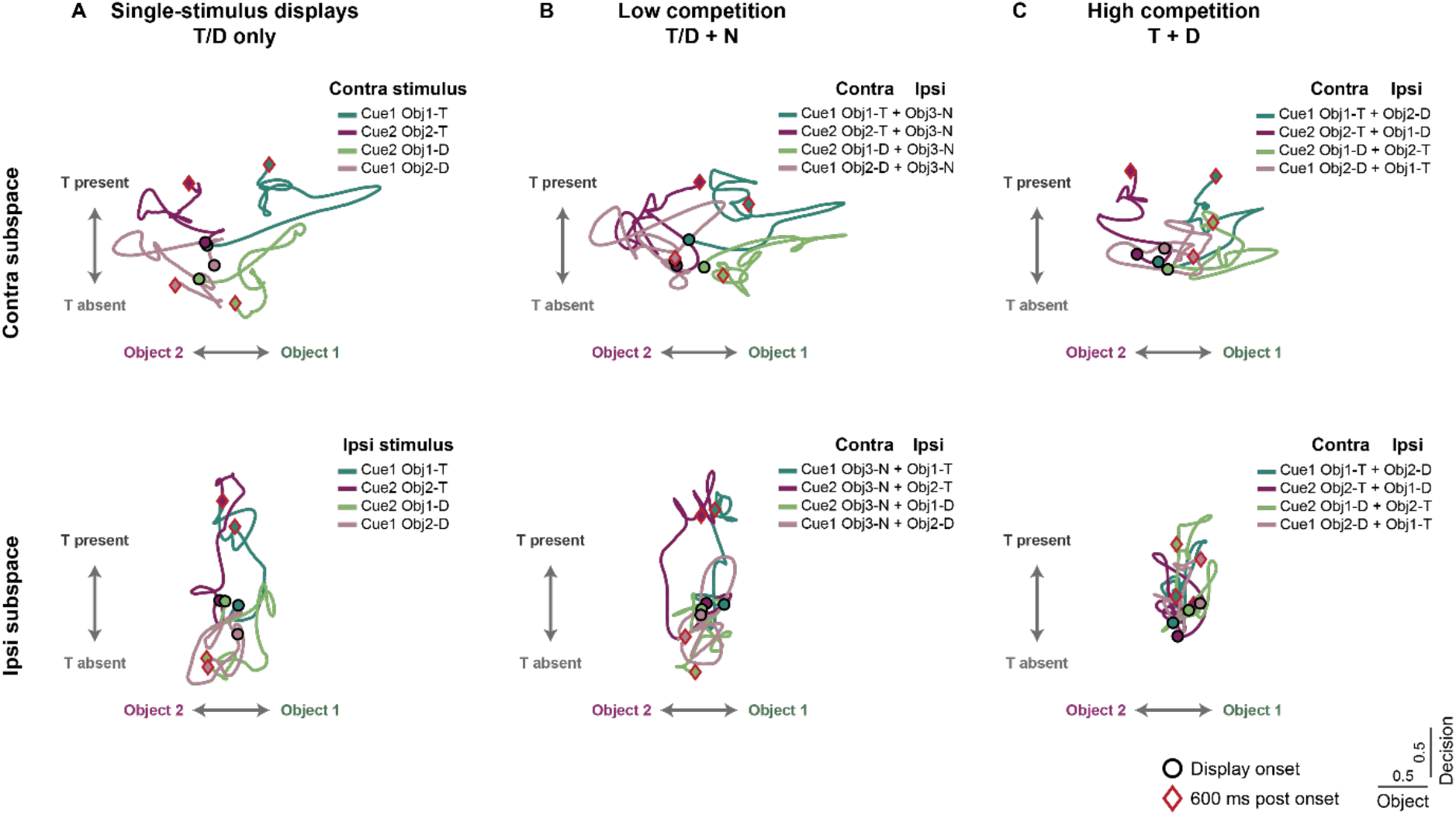
Dynamics of the neural state space. **A:** Single-stimulus displays. Population activity is projected onto the object and decision components for contralateral (top) and ipsilateral (bottom) stimuli, each projected onto their own subspace. Data are the same as in Figures 3A and 4A, top left and bottom right panels, shown here as trajectories in the state space. Cross-validated trajectories are shown as averages of the two halves of the data projected onto the subspace of the other half of the data. Responses are smoothed by a ±25 ms window and are shown from stimulus onset (black circle) to 600 ms post stimulus onset (red diamond). Scale for stimulus and decision responses shows normalized firing rate (sp/s). **B:** Low competition. Data are the same as in Figures 3B and 4B, top left and bottom right panels. **C:** High competition. Data are the same as in Figure 6A and 6B.

### Distributed, mixed and independent task-related representations

The neuronal weights associated with each axis of a subspace reflect the strength of contribution of each neuron’s activity to the overall population coding, as well as its preference depending on the sign (e.g., object 1 or 2 for object coding). For each axis, the distribution of weights across neurons was unimodal and centered around 0 (Figure 8A), demonstrating that coding of task-relevant variables was distributed with varying levels of contributions across neurons, rather than driven by a small subpopulation. Weights on each axis were correlated across contralateral and ipsilateral subspaces (Figure 8A), with the largest correlation for the decision component where coding was relatively strong in both contralateral and ipsilateral subspaces. Correlations were significant, however, even for object and cue, despite weak coding of object in the ipsilateral subspace and cue in both subspaces. These correlations demonstrate that the representations in the two hemispaces are related. Finally, a key question is whether coding across the neural population of different task variables is driven by separate sub-populations of neurons, or whether computation is done within the same neural circuit for all task-relevant variables. We addressed this question by correlating the rectified neuronal weights of pairs of task variables (Figure 8B). Weights were rectified in order to consider the magnitude of contribution of each neuron rather than its preference (e.g to object 1 or 2). Correlations were generally low, except for a positive correlation between decision and object weights in the ipsilateral subspace. The result shows the dominance of mixed selectivity, with a neuron’s contribution to coding of each task variable rather independent of its contribution to others.

**Figure 8:**
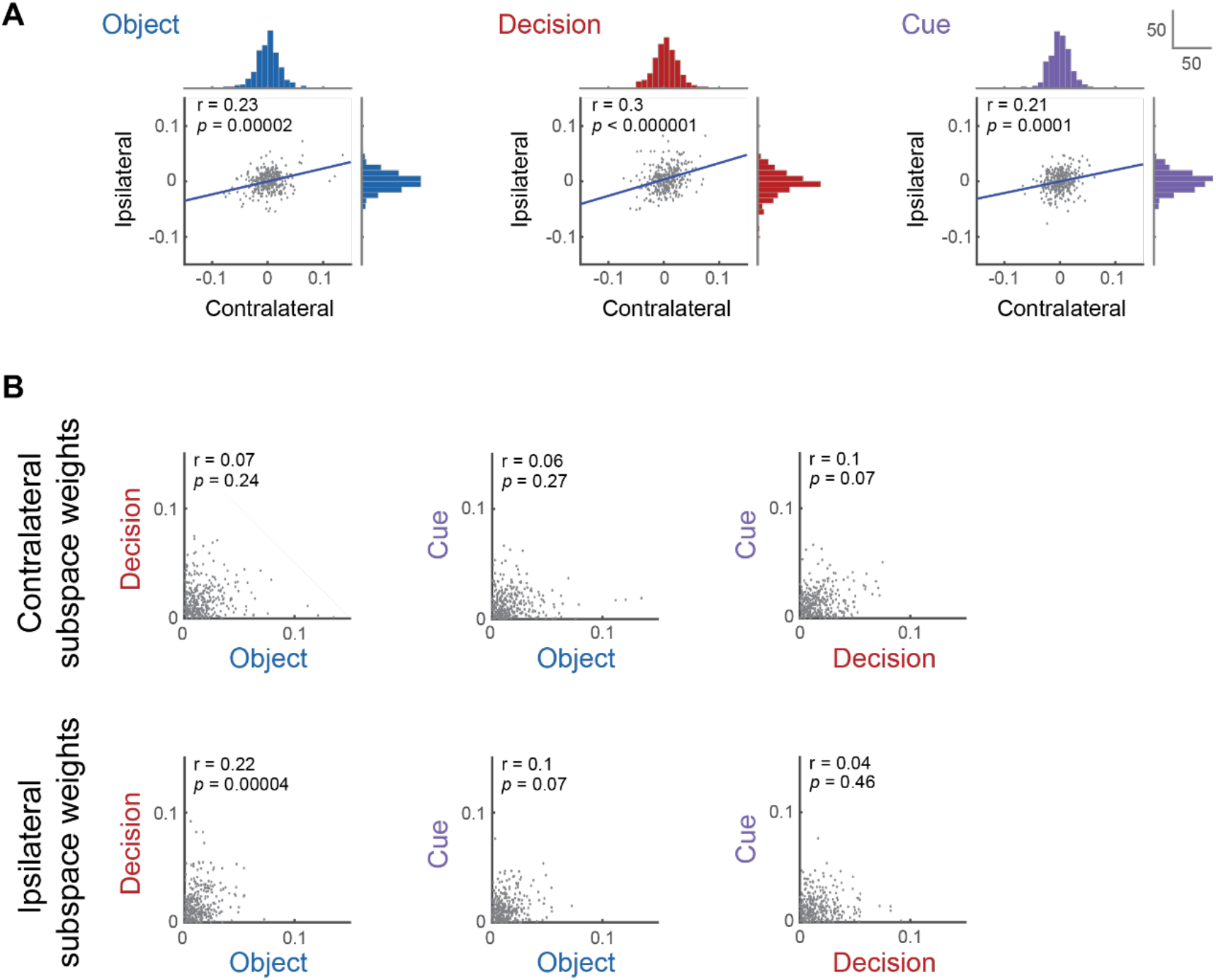
Distributed, mixed and independent task-related representations across the neuronal population. **A:** Relationship of weights in contralateral and ipsilateral subspaces for each task variable (subspace axis). Each dot is one neuron. Pearson correlation coefficient, *p* value and a regression line are indicated within each plot. Histograms at top (contralateral subspace) and right (ipsilateral subspace) show weight distribution across neurons, with y axis scale on the top right. **B:** Rectified weights of individual neurons for each pair of axes, for the contralateral (top) and ipsilateral (bottom) subspaces. Absolute weights are plotted to show strength of selectivity regardless of preference. Spearman’s rank correlation coefficient and *p* value are indicated within each plot. For all plots, weights of each neuron and axis are averaged across the subspaces derived from the two halves of the data.

### Reliability

We constructed the low-dimensional subspaces by using a split-half cross-validated approach. To investigate the reliability of representations, we correlated weights derived from each half of the data (Figure 9). Reliabilities were highly significant, particularly for the components that showed strong coding: the decision variable in both hemifields (r = 0.43 and r = 0.53 for contralateral and ipsilateral subspaces, respectively, both *p* < 0.000001) and the object variable in the contralateral subspace (r = 0.49, *p* < 0.000001). Even for other variables, however, these data confirm some stability in coding in each subspace.

**Figure 9:**
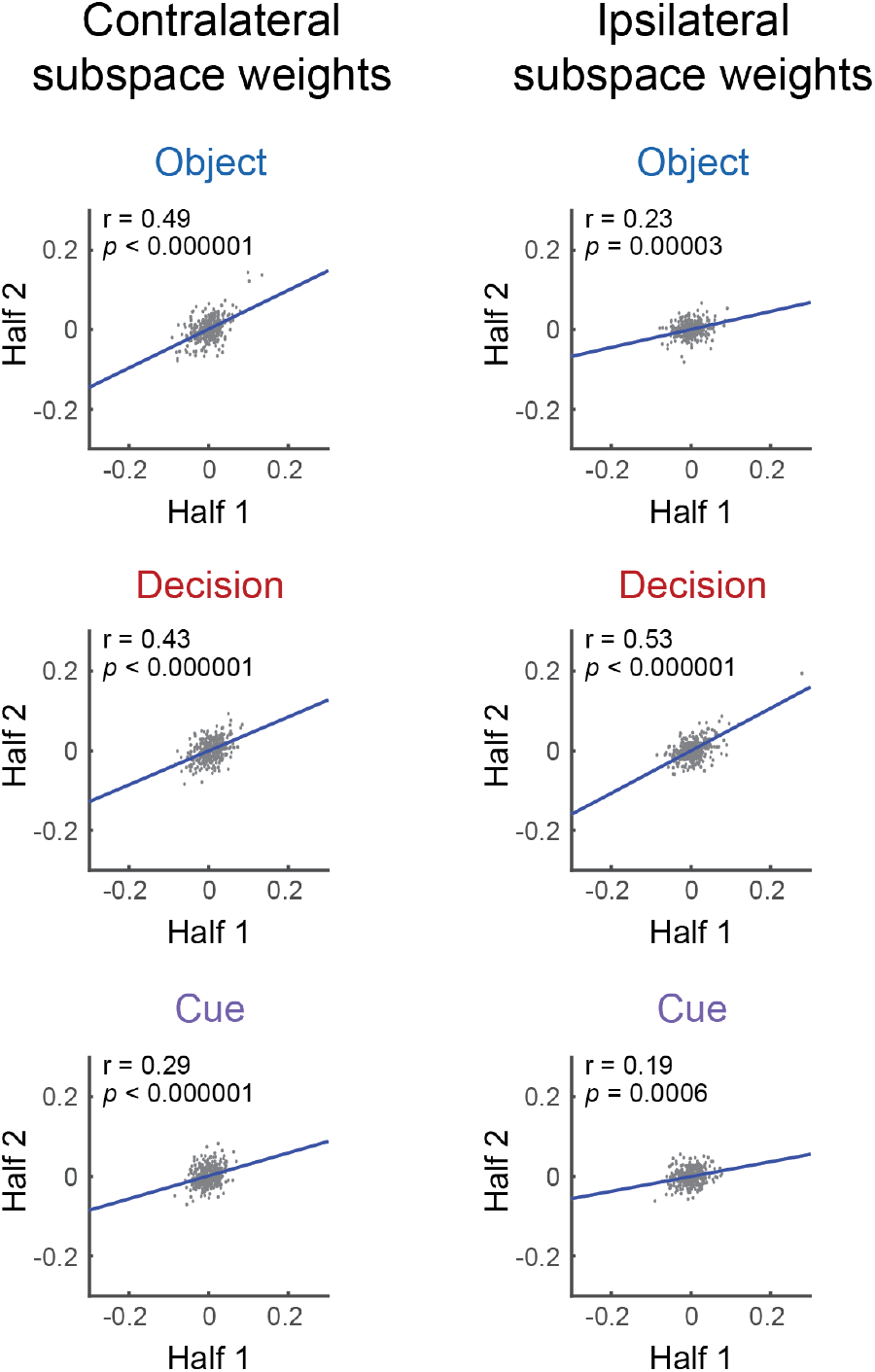
Reliability of coding in each subspace. For each task variable (subspace axis), plots show correlation between weights derived from separate halves of the data. Each dot is one neuron. Pearson correlation coefficient, *p* value and a regression line are indicated within each plot.

## DISCUSSION

In this study we investigated prefrontal coding during a context-dependent decision-making task while items compete for attention. We used a data set from our previous work where we showed how activity in a prefrontal cell population shifts from dominance by a contralateral stimulus to dominance by a behaviorally relevant target item (26). Here, we use dPCA to construct a task-related low-dimensional subspace that reflects the learned task structure and examine separate coding of cue, object and decision and their dynamics throughout the course of a trial. We then track coding of competing items along the same subspace and show how task-related codes evolve and integrate while competition resolves.

Our results demonstrate variable temporal profiles for coding of cue, object and decision in the prefrontal cortex. Prefrontal coding of task-related information has been previously demonstrated across a wide range of variables, including visual properties (e.g., motion color), tactile input, cues, reward value, belief updating and decision (27,29,38–42). Here, we emphasize not only how coding of task-related variables dynamically evolves but also how they integrate to implement a control program of the task as reflected in the extracted multidimensional subspace. This same subspace can be then used to investigate coding in an extension of the task in which items compete for attention, unravelling neural representations as competition unfolds.

Our results suggest that communication between hemispheres plays an integral role in attentional control. We illustrate the task-related subspace and flow of information within and between hemispheres in the model in Figure 10. In the task-related subspaces for single stimulus displays (Figure 10A), coding along the cue and object axes (top) as well as the decision axes (bottom) is shown for the two hemispheres relative to the stimulus presentation hemifield. While we recorded activity in a single hemisphere for contralateral and ipsilateral stimuli, in the model, findings are re-cast to show inferred activity in each hemisphere for a single stimulus. Arrows between top and bottom sections show movements along the decision axis (bottom) following from activity at different loci within the object-cue subspace (top). Extension to an example high-competition display is shown in Figure 10B.

**Figure 10.**
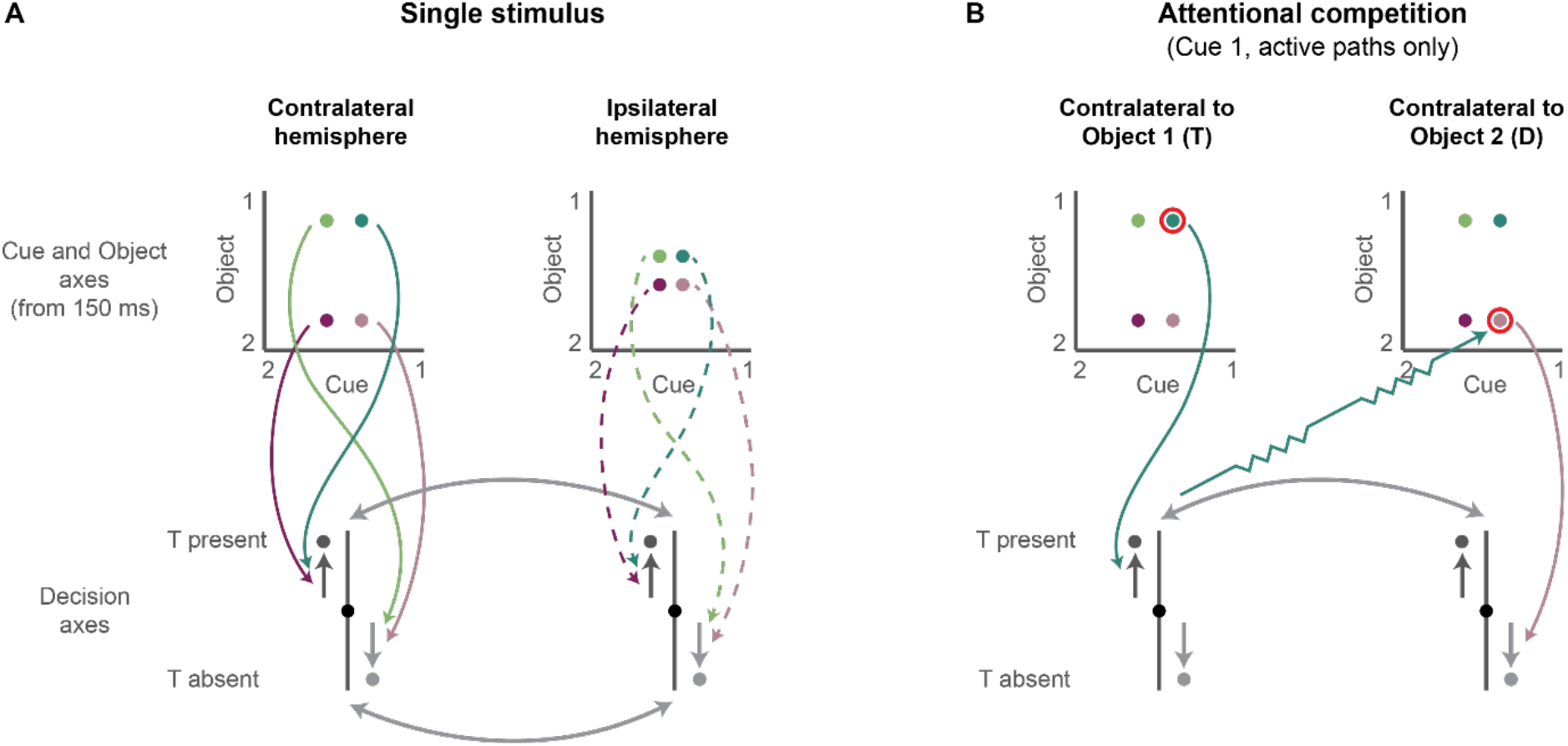
A model for attentional competition in prefrontal cortex. **A. Single-stimulus displays.** The task subspace for cue and object axes (top) and the decision axis (bottom) is shown for the two hemispheres relative to the stimulus presentation hemifield. Each colored dot shows representation of object and cue information for each of the four single-stimulus conditions. Shortly after stimulus onset, strong coding emerges in the hemisphere contralateral to the stimulus. Only weak object and cue coding are seen in the ipsilateral hemisphere. Integrated object and cue information then drives coding along the decision axis, mainly in the contralateral hemisphere (colored solid arrows), and only weakly in the ipsilateral hemisphere (colored dotted arrows). Coding along the decision axis moves towards the behavioral decision (dark and pale gray arrows beside decision axis), and as it develops is quickly transferred between hemispheres (pale gray arrows between the decision axes). **B. Attentional competition.** The task subspaces are shown for competition of a target (T) and distractor (D), with arrangement similar to the single-stimulus displays. For simplicity, within each hemisphere, partially-overlapping decision axes for contralateral and ipsilateral stimuli are collapsed into one (see Discussion). In this example trial, cue 1 sets object 1 as target and object 2 as distractor (red circles). Subspaces in each hemisphere are shown relative to the contralateral stimulus, with colored arrows showing active paths of influences driven by this example stimulus. Initially, object coding in each hemisphere is driven by the contralateral stimulus (top). Integration of cue and object then tries to drive the decision state in each hemisphere towards the corresponding behavioral decision (cyan and pink arrows). As the decision state approaches ‘target present’ in one hemisphere, however, this suppresses representation of the distractor object in the opposite hemisphere (zigzag arrow). Suppression may be direct or indirect (see Discussion). Finally, the ‘target present’ decision from the hemisphere contralateral to the target is transferred to the other hemisphere, though only slowly and weakly owing to that hemisphere’s initial processing of the distractor.

For a single stimulus (Figure 10A), object coding in the contralateral hemisphere rose to a peak around 150 ms from display onset, followed by sustained coding throughout the rest of the trial. Cue coding in this hemisphere was weak, but followed a generally similar time course. Figure 10A illustrates the locus of activity in the object-cue subspace for the relatively stable period from 150 ms onwards, with colored dots for the four single-stimulus conditions in our task. Integrated object and cue information then drives coding along the decision axis (solid colored arrows) towards the correct behavioral decision, either ‘target present’ or ‘target absent’ (dark and pale gray arrows, respectively), with a gradually increasing strength of coding over time. These coding patterns reflect context-dependent computation in the prefrontal circuit dominated by the contralateral stimulus, in line with previous reports (26,27,33,36).

In contrast, in the ipsilateral hemisphere there was only weak object and cue coding, as illustrated with the four colored dots close to the center of the two axes in Figure 10A (top right). Object coding for a single ipsilateral stimulus, although weak, was sustained for a time window similar to the peak of coding for a contralateral stimulus, consistent with our previous report for the same dataset where a small proportion of cells were selective for an ipsilateral object identity (36). Importantly, despite the weak object and cue coding on the ipsilateral hemisphere, which can have only limited effect on the decision (dotted colored arrows), a strong decision state developed, similar in strength and dynamics to the one on the contralateral hemisphere. Given weak object and cue information in the ipsilateral hemisphere, a likely mechanism to drive a strong decision trajectory is cross-hemisphere transfer of decision information from the opposite hemisphere (pale gray bi-directional arrows between the decision axes). Bi-directional exchange of decision information between the hemispheres then further reinforces the development of a coherent decision state in both hemispheres, either ‘target present’ or ‘target absent’.

Within each hemisphere, moderate but highly significant correlations between contralateral and ipsilateral dPCA weights (Figure 8A) revealed some overlapping, though not identical, coding of each task variable for contralateral and ipsilateral stimuli. The same point is reflected in the cross-subspace projections (top right and bottom left plots in figures 3–5). Especially important is the decision axis, which in each hemisphere showed strong coding for both contralateral and ipsilateral stimuli. Partial overlap of the two axes suggests an integrated ‘target present’ or ‘target absent’ decision. Partial separation likely reflects the different responses (saccade directions) required for contralateral and ipsilateral stimuli. The absence of saccadic activity in this cell population (26) suggests an abstract response decision rather than specific motor preparation. Mostly negligible weight correlations across different task variables (Figure 8B) demonstrate mixed selectively of prefrontal neurons (8), such that a neuron’s contribution to any one representational axis was largely independent of its contribution to others. Notably absent were negative correlations, which might be expected if dedicated neural populations coded object and cue, then feeding into a further dedicated population for decision. Instead, overlapping populations could contribute to coding each task variable (43), as would be expected, for example, in a recurrent network (24,27).

A primary focus in this study was the dynamics within the task-related state space when items compete for attention. The importance of levels of competition to behavioural outcome is well established in human visual search experiments (31,44). Items that are not currently targets but have been frequently experienced as targets throughout learning history impose a large conflict and compete strongly with target items for attention. In contrast, less conflict is evoked by items that can always be ignored. A similar manipulation of competition was applied in our study, with distractor objects strongly competing for attention, and therefore expected to introduce a substantial effect on coding within the state space, and neutral items posing only little competition with potentially little effect on coding.

Indeed, adding a low-competition neutral item to a target or distractor had very little effect on coding of object, decision and cue information. One striking effect, however, is that object coding in the ipsilateral hemisphere – already weak for a single stimulus – was now completely eliminated by the accompanying neutral item in the opposite hemifield (Figure 3). Despite the absence of object coding in the ipsilateral hemisphere, strong decision coding still developed (Figure 4). This finding adds weight to our proposal that the decision is first computed in the contralateral hemisphere, then communicated from one hemisphere to the other.

A different picture was observed for displays containing both a target and distractor, when attentional competition was high. Movements in the representational space for this case are illustrated in Figure 10B, for an example display with cue 1 followed by a target in one hemisphere and a distractor in the other. Note that, within each hemisphere, there are potential movements along two decision axes, one for the contralateral and one for the ipsilateral stimulus, but given the partial overlap of these two axes, here for simplicity they have been collapsed into one. Figure 10B shows arrows just for the active paths, i.e., those driven by the particular combination of stimuli present in the example display.

The results show that, in each hemisphere, object coding was dominated by the contralateral stimulus, either a target or distractor (Figure 6A). Given the connections inferred for the single stimulus case (Figure 10A), these two separate object representations should attempt to drive decision coding in the two hemispheres in opposite directions (cyan and pink arrows). For the hemisphere contralateral to the target, decision activity evolved as expected (Figure 6B), but for the hemisphere contralateral to the distractor, a very different picture emerged. First, the data suggest a rapid shut-off of the distractor object representation (Figure 6A). Then decision coding moves, not towards the ‘target absent’ decision, but slowly and weakly towards the (task-appropriate) ‘target present’ decision. To achieve this outcome, Figure 10B shows suppression from the ‘target present’ end of the decision axis in one hemisphere to the object representation in the opposite hemisphere (zigzag arrow). As this suppression develops, object representation is lost in the hemisphere contralateral to the distractor, leaving the ‘target present’ decision to be transferred from the opposite hemisphere. Though the zigzag arrow in Figure 10B suggests direct interaction between the two frontal lobes, this is only one possible way for the ‘target present’ decision in one hemisphere to suppress object information in the other. A different possibility would be for detection of the task-relevant target to feed back to earlier cortical levels, biasing competition to this target and suppressing the accompanying visual representation of the distractor. In either case, as the logic of the task requires, the object that should control the decision comes finally to dominate neural activity of both hemispheres along the decision axis, though in the hemisphere contralateral to the distractor, this dominance is achieved only slowly and incompletely, presumably reflecting a remaining influence from initial distractor processing.

These results match and further strengthen the findings reported by Kadohisa et al. (26), where representation in the high-dimensional space for an ipsilateral target remained unaffected when a contralateral neutral item was added, but showed a large divergence when a contralateral distractor was added. They are also consistent with attentional competition in visual areas, where early sensory-driven responses are followed by responses dominated by the most behaviorally relevant stimulus (30,45). While we focus here on competition for visual attention, similar principles of cross-hemisphere exchange of information would likely play a central role in prefrontal coding for goal-directed behavioral decisions more generally.

Figure 10 exemplifies the kind of computational structure required for control of complex behavior. In prefrontal cortex, such structures must be assembled by learning; for example, learning that cue and object information must be combined to determine oculomotor choices, and that the cued target should dominate in a two-stimulus display. In the broader context, this is an example of just one task that can be learned and coded in the prefrontal cortex. Modeling studies have demonstrated how such computation can be reproduced within a neural network (27,46). Furthermore, one network can implement many such computations for different tasks that are learned through experience, creating a task space from which task representations can be composed to facilitate learning of new task structures (47). We demonstrate here an implementation of one building block within such task space and how it can be used for neural computation in an extended task with competing items.

Dimensionality reduction techniques such as dPCA offer a window onto such learned computational structures and their dynamics within the subspace. Our results highlight integration of cue and stimulus information while a focused attentional state develops, with distinct temporal profiles of coding for these task variables, as well as communication between hemispheres as key for contextdependent processing of competing items. The results provide a comprehensive and detailed account of information coding in the prefrontal cortex as a learned decision unfolds and attentional competition resolves.

## MATERIALS AND METHODS

### Subjects

Subjects were two male rhesus monkeys (*Macaca mulatta*) weighing 11 (monkey A) and 10 kg (monkey B). All the experimental procedures were conducted in accordance with the Animals (Scientific Procedures) Act 1986 of the UK. All the procedures were in compliance with the guidelines of the European Community for the care and use of laboratory animals (EUVD, European Union directive 86/609/EEC) and were licensed by a Home Office Project License obtained after review by Oxford University’s Animal Care and Ethical Review committee.

### Task

A cued target detection task was used (Figure 1A). Based on training prior to recording, two cues (cue 1, cue 2) were associated with one target object each (object 1, object 2), and a third neutral object (object 3) was not associated with any cue. Figure 1A shows the cue-target pairs for monkey A; different items were used for monkey B. A red dot at the center of the screen marked the start of a trial; the monkey was required to fixate throughout the trial until the saccadic response at the end (fixation window: 5° x 5° for monkey A, 4° x 4° for monkey B). Following fixation for 1,000 ms, a cue stimulus (2° x 2°) was presented at the center of the screen for 500 ms, indicating the target for the current trial, followed by a randomly varying delay (400-600 ms for monkey A and 400-800 ms for monkey B). Next, a choice display was presented (500 ms), including either a single object or two objects. Objects (2° x 2°) were always centered on the horizontal meridian. In single-stimulus displays, the object was presented 6° to the left or right of fixation, determined randomly. The position of the objects was therefore either contralateral or ipsilateral hemifield to the recorded hemisphere. The object could be either a target (T) if associated with the preceding cue, a distractor (D) if associated with the other cue, or a neutral stimulus (N) if not associated with any cue and therefore never a target (object 3). To test for attentional competition, two-stimulus displays contained one stimulus in each hemifield, with combinations of target and neutral objects (T + N, low competition), target and distractor objects (T + D, high competition), and distractor and neutral objects (D + N, target absent trials). Objects were presented 6° to the left and right of fixation, and the right-left or left-right configuration was randomly determined. To avoid response bias, the frequencies of the choice display types were adjusted to include a target object in half of all single-stimulus and half of all two-stimulus displays. The frequencies of the main display types were otherwise the same. A trial was terminated without reward if there was a premature saccadic response outside the fixation window during the trial, and these trials were excluded from the analysis.

The choice display was followed by an additional delay period (randomly varying, 100-150 ms for monkey A, 300-500 ms for monkey B). The fixation dot then turned green, indicating the go signal and response interval. For trials that included a target in the display (target present), the monkey had to saccade to the remembered T location (target window: 6° x 6° for monkey A, 3.5° x 3.5° for monkey B) and was rewarded immediately for a correct saccade with a drop of liquid. For trials that did not include a target in the display (target absent), the monkey had to fixate for the whole response interval (1,000 ms) and was then rewarded either for a further saccade (monkey A) or immediately (monkey B).

For monkey A, cues varied randomly between trials in some sessions, while other sessions included alternating short blocks of fixed cues (15-20 trials per block). Physiological data were very similar in the two cases and were combined. For monkey B, cues always varied randomly between trials.

Overall, the task included six main choice display types (three single-stimulus and three two-stimulus displays), and a total of 24 conditions defined by combinations of cue, display type and hemifield. Four neutral-only conditions were not included in the analysis, along with a small number of D + D trials present in some sessions.

### Recordings

Each monkey was implanted with a custom-designed titanium head holder and recording chamber(s) (Max Planck Institute for Biological Cybernetics, Tübingen, Germany), fixed on the skull with stainless steel screws. Chambers were placed over the lateral prefrontal cortex of the left (AP = 25.3, ML = −20.0; AP, anterior-posterior; ML, medio-lateral) and right (AP = 31.5, ML = 22.5) hemispheres for monkey A and the right hemisphere (AP = 30.0, ML = 24.0) for monkey B. Recording locations for each animal are shown in Figure 1B. Under each chamber, a craniotomy was made for physiological recording. All surgical procedures were aseptic and carried out under general anesthesia. We used arrays of tungsten microelectrodes (FHC, Bowdoin, ME, USA) mounted on a grid (Crist Instrument, Hagerstown, MD, USA) with 1 mm spacing between adjacent locations inside the recording chamber. The electrodes were independently controlled by a hydraulic, digitally controlled microdrive (Electrodes Drive, NAN Instruments, Nof Hagalil, Israel, for monkey A; Multidrive 8 Channel System, FHC for monkey B). Neural activity was amplified, filtered, and stored for offline cluster separation and analysis with the Plexon MAP system (Plexon Inc, Dallas, Texas, USA). Eye position was sampled using an infrared eye tracking system (120 Hz, ASL, Bedford, MA, USA, for monkey A; 60 Hz, Iscan, Woburn, MA, USA, for monkey B) and stored for offline analysis. Data were recorded over a total of 140 daily sessions. Before starting the task, microelectrodes were advanced until neuronal activity could be isolated. Neurons were not preselected for task-related responses. At the end of the experiments, animals were deeply anaesthetized with barbiturate and then perfused through the heart with heparinized saline followed by 10% formaldehyde in saline. The brains were removed for histology and recording locations were confirmed on dorsal and ventral frontal convexities and within the principal sulcus.

### Data and analysis

Only data from successfully completed trials were analyzed, and only units with a minimum of four trials in each condition were included in the analysis. All statistical analyses were conducted using MATLAB (MathWorks Inc.). We used customized code for the analysis and the demixed PCA toolbox (28) to construct the low-dimensionality subspaces.

For all analyses, spike data of each unit were first smoothed with a Gaussian kernel of SD=20 ms, cutoffs ±1.5 SD. The analysis focused on the choice display epoch of a trial where cue and stimulus information are combined to reach a decision. We therefore used data from each trial from −200 ms to 600 ms from choice display onset, throughout which the animals kept fixation. Data for each neuron were normalized across all trials, time points and conditions by subtracting the mean and dividing by SD prior to any analysis.

### Task-relevant low-dimensional neural state space

The demixed principal component analysis (dPCA) is described in detail in Kobak et al. (28). Briefly, it decomposes the high dimensional space of population activity into a small number of task-related components while minimizing the error between the reconstructed signal and average activity for each level of the different task variables. The resulting compressed subspace both captures the majority of variance in the data, as well as keeps the task-related components demixed, such that each component captures variance that is related mostly to one task variable only and the levels of this variable are separable to allow decodability.

We constructed subspaces based on the single-stimulus T and D displays. Since population activity in the prefrontal cortex is highly affected by hemifield (26,30,34–37), we constructed separate subspaces for the contralateral and ipsilateral hemifields. We extracted separate components for object identity [(cue 1, object 1 + cue 2, object 1) – (cue 1, object 2 + cue 2, object 2)], decision [(cue 1, object 1 + cue 2, object 2) – (cue 1, object 2 + cue 2, object 1)] and cue ((cue 1, object 1 + cue 1, object 2) – (cue 2, object 1 + cue 2, object 1))). We used the first component of each task variable, which explains the largest amount of variance (see Results). Calculation of explained variance by each of the dPCA components was conducted using the dPCA toolbox and is described in detail in Kobak et al. (28). Briefly, explained variance was computed in a standard way as the fraction of variance explained in peri-stimulus time histograms (PSTHs), subtracting the unexplained variance (difference between the PSTHs and the reconstructed signal) from the total variance of the PSTHs and dividing by the total. Using the PSTHs rather than single trial data was required because data were sequentially recorded in sessions over different days. The explained variance was further decomposed into the contribution of the different task variables by using the marginalized PSTH across these variables.

### Population coding of task-relevant information

We tracked the dynamics of cue, object and decision information within the task-related subspace and how it evolves when two items compete for attention. To investigate these temporal trajectories of representations of task-relevant information, we projected the data onto the compressed subspaces. To ensure the generalizability of the compressed subspaces and avoid over-fitting, we employed a split-half cross-validation approach. We split the data for each neuron and condition into two halves (odd and even trials). Within each half, data for each condition were averaged across trials in 1 ms bins to create a PSTH. We then computed separate subspaces for each half, and projected data from each half onto the subspace constructed from the other half. Results from these two crossvalidated projections were then averaged. Signs of neuronal weights for each component were reversed when required to ensure that all the four subspaces (two halves, two hemifields) were compatible.

### Statistical analyses

To measure coding of task-relevant information across the subspace, we subtracted the averaged projections of one level of each task variable on the corresponding component from the averaged projections of the other level. For example, for object information, the average projection of object 2 (across the two cues/decisions) on the object component was subtracted from the average projections of object 1 (across the two cues/decision). Similar measures were obtained for decision information (T present – T absent) and cue information (cue 1 – cue 2). Information was computed for each time point (1 ms bins) and averaged across subspaces for the two halves of the data.

We then used permutation analysis to determine statistical significance of these information measures, with cluster size correction for multiple comparisons across time points. For each comparison, a null distribution was generated by permuting condition labels for the trials within each half of the data (odd and even trials) then repeating exactly the same procedure as the main test: computing PSTHs, projecting the PSTHs onto the relevant component in the other half subspace, computing the difference using the appropriate contrast to measure coding and averaging across halves. This was repeated 1000 times. To estimate the distribution of cluster sizes expected by chance, we used a leave-one-iteration out approach. For each iteration, we marked time points at which the measured difference was greater than the value in all remaining iterations, then selected the cluster with the maximal number of such time points. To establish significance in the real data, we first marked time points at which the measured difference in the real data was greater than the values in all the permuted data (i.e., α = 0.001, one-tailed) and considered as significant only clusters larger than all maximum clusters in the permuted data (i.e., α = 0.001, one-tailed). Permutation analysis to compare information in two-stimulus and single-stimulus displays was done in a similar way, permuting condition labels between the appropriate conditions for each comparison. Because the subspaces and statistical tests were done using split-half cross-validation, comparisons between any two or more conditions were not prone to biases depending on whether they were used to construct the subspace (i.e., single-stimulus displays) or not (i.e., two-stimulus displays).

### Correlations of neuronal weights

We used correlations across the neural population to test for dependence between neuronal dPCA encoder weights between subspaces and axes. For dependence between axes (objects, decision, and cue), we averaged the components weights across the subspaces of the two halves of the data, rectified them, and correlated all three possible pairs of axes using Spearman’s rank correlation, for each hemifield subspace separately. The weights were rectified because the measure of interest for this analysis is the relative contribution of each neuron as reflected in the absolute weight, rather than its preference as reflected in the sign (e.g., preference for object 1 or object 2). To test for the dependence of representations across the contralateral and ipsilateral subspaces, for each subspace we averaged weights from the two halves of the data, and Pearson-correlated them across the two hemifields. Similarly, to test for the reliability of each set of neuronal weights within a subspace, we used Pearson’s correlation between weights extracted from the two independent halves of the data.

## DATA AND CODE AVAILABILITY

The raw neuronal data and custom analysis scripts that support the findings in this study will be made freely available for download at a public repository upon publication. The code for the dPCA toolbox is freely available for download (28).

## ACKNOWLEDGMENTS

This work was supported by MRC intramural programme SUAG/002/RG91365, a research award 220020081 from the James S. McDonnell Foundation, and grant 101092/Z/13/Z from the Wellcome Trust. Y.E. was supported by a Royal Society Dorothy Hodgkin Research Fellowship DH130100. We thank Daniel Mitchell for useful comments and advice throughout this study.

## AUTHOR CONTRIBUTIONS

Y.E. conceived the study, analyzed data and wrote the manuscript. M.Ka., P.P. and M.Ku. collected data. M.Ka., P.P., N.S., M.B., M.Ku and J.D. contributed to original task design and project administration. J.D. conceived the study, supervised all aspects of the project and wrote the manuscript.

